# Integrative proteogenomics using ProteomeGenerator2

**DOI:** 10.1101/2023.01.04.522774

**Authors:** Nathaniel Kwok, Zita Aretz, Sumiko Takao, Zheng Ser, Paolo Cifani, Alex Kentsis

## Abstract

Recent advances in nucleic acid sequencing now permit rapid and genome-scale analysis of genetic variation and transcription, enabling population-scale studies of human biology, disease, and diverse organisms. Likewise, advances in mass spectrometry proteomics now permit highly sensitive and accurate studies of protein expression at the whole proteome-scale. However, most proteomic studies rely on consensus databases to match spectra to peptide and proteins sequences, and thus remain limited to the analysis of canonical protein sequences. Here, we develop ProteomeGenerator2 (PG2), based on the scalable and modular ProteomeGenerator framework. PG2 integrates genome and transcriptome sequencing to incorporate protein variants containing amino acid substitutions, insertions, and deletions, as well as non-canonical reading frames, exons, and other variants caused by genomic and transcriptomic variation. We benchmarked PG2 using synthetic data and genomic, transcriptomic, and proteomic analysis of human leukemia cells. PG2 can be integrated with current and emerging sequencing technologies, assemblers, variant callers, and mass spectral analysis algorithms, and is available open-source from https://github.com/kentsisresearchgroup/ProteomeGenerator2.

## Introduction

Mass spectrometry-based proteomics now enables genome-scale measurements of proteomes from diverse cells, tissues, and organisms. Most commonly, this approach leverages high-resolution mass spectrometric measurements of proteolyzed proteins, combined with tandem peptide fragmentation to match observed spectra with those expected from their amino acid composition. Current approaches for peptide and protein identification use advanced statistical and graphical methods to match mass spectra and estimate their confidence.^1,2,3,4,5,6,7^ Recent tools including MSFragger^6^, MetaMorpheus^8^, TagGraph^7^, and SAMPEI^9^ also permit agnostic discovery of modified peptides and proteins, including amino acid substitutions and chemical modifications. With the exception of completely *de novo* mass spectrometry identification algorithms which currently lack the requisite accuracy and efficiency for genome-scale studies, all current methods depend on matching observed mass spectra to reference proteome databases.

However, recent studies in many organisms and in humans have revealed significant protein sequence variation due to the presence of somatically acquired genetic variants, alternative transcription, and mRNA splicing, which are not necessarily annotated in reference databases. For example, most human tissues in healthy individuals acquire somatic nucleotide substitutions, insertions, deletions and DNA rearrangements, leading to the production of variant protein isoforms.^10^ Similarly, in recent studies, at least one third of genes in diverse organisms can exhibit alternative transcription, leading to the production of N-terminally extended proteins or alternative reading frames.^11^ And human diseases, including cancer in particular, tend to be defined by the presence of proteins with altered and pathogenic sequences.^12^ Thus, there is a pressing need for improved methods for the discovery of non-canonical and sample-specific proteomes.

Several approaches have been developed to generate sample-specific mass spectrometry search databases using RNA sequencing.^13–16^ However, these methods depend on reference transcriptome annotation and cannot leverage recent advances in strand-specific or long-read RNA sequencing. Recently, to enable facile and scalable proteogenomics, Cifani et al and Cesnik et al leveraged the snakemake extension language to develop the ProteomeGenerator and Spritz software tools, respectively.^17,18^ Snakemake^19^ is a programming workflow management system, which uses simple commands accessible to both expert and novice users, to generate and execute Python programs, while maintaining correct dependencies and enabling scalable development of complex modular computational methods, as required for comprehensive proteogenomics. In particular, Spritz used the hisat2^20^ read aligner and GATK variant caller^21^ to identify nucleotide variants, such as substitutions, insertions and deletions present in expressed transcripts, leading to their inclusion in sample-specific proteome databases for mass spectrometry analysis. ProteomeGenerator leveraged recently developed methods for transcriptome assembly (StringTie^22^ and TransDecoder^23^) to capture proteome variation induced by alternative transcription, splicing, and genome rearrangements.

Here, we extend the ProteomeGenerator framework to develop ProteomeGenerator2 (PG2). PG2 integrates genome DNA sequencing and variant calling with referenced and de novo transcriptome assembly to enable the identification of genetic and mutational variants from nucleotide substitutions, insertions, and deletions, as well as non-canonical protein isoforms arising from alternative transcriptional start sites, intron retention, cryptic exon splicing, and gene fusions. Integral to the complete workflow of PG2 is the creation and usage of sample-specific genome assemblies (via bcftools^24^), which are generated from any combinations of genomic sequencing and/or structural variation in tandem with a reference genome. We demonstrate that a sample-specific genome-based approach not only enables generation of precise, comprehensive, and extensible proteome databases, but also improves the accuracy of RNA read mapping and transcriptome assembly.

## Materials and Methods

### ProteomeGenerator2 DNA sequence analysis using whole-genome, whole-exome, or target gene sequencing

#### Whole-genome sequencing alignment

Paired-end whole-genome sequencing read FASTQ files were trimmed for adapter sequences with trimmomatic (v0.39).^25^ The trimmed FASTQ files were converted to unmapped BAM, sorted by read name, and then aligned to the reference genome with *BWA mem*. The resulting mapped BAM was then merged with the original unmapped BAM with *picard MergeBamAlignment*^*26*^ for maximal data utilization.

#### Library merging, duplicate marking, and base score recalibration

Readgroup BAM files were merged with *picard* (v2.18.21) *MergeSamFiles*. Duplicates were marked with *gatk*^*27*^ (v4.1.4.1) *MarkDuplicates*, coordinate-sorted with *gatk SortSam*, and indexed with *samtools*^*24*^ *index*. Base recalibration was performed with *gatk BaseRecalibrator*, with hapmap v3.3 (‘HapMap’) and Mills and 1000G gold standard indels (‘Mills’) (as available in the v0 GATK resource bundle for hg38) used for known SNPs and indels. The recalibrated base reports were applied to both mapped and unmapped reads with *gatk ApplyBQSR*. The recalibrated BAMs were gathered and combined with *picard MergeSamFiles*.

### Genome variant calling

#### Germline short indel/genotyping with GATK4 HaplotypeCaller

Germline variants were called on prepared BAM files with *gatk* (v4.1.4.1) *HaplotypeCaller*^*21*^, as specified by the ‘WGS hg38 calling regions’ provided by the GATK resource bundle. For datasets with matched samples, each tumor/normal or experiment/control was processed individually. Variant calling was scattered over 50 intervals. Called variant gVCFs were genotyped with *gatk GenotypeGVCFs*. Genotyped VCFs and prepared BAM files were deployed to establish a 2D convolutional neural network variant scoring model with *gatk CNNScoreVariants*. A *Singularity*^*28*^ container was utilized for *TensorFlow*^*29*^ support. Score tranches were assigned using HapMap and Mills as snp and indel resources, respectively, with tranche cutoffs set at 99.9 and 96.0 for snps and indels, respectively. Variants that were not assigned a “PASS” were filtered out with *bcftools* (v1.10.2) *view*. Interval-wise VCFs were merged with *picard* (v.2.18.21) *MergeVcfs*. In-cohort samples were prepared for consolidation by consolidating their respective names with *bcftools reheader*. Finally, in-cohort sample VCFs were consolidated and de-duplicated with *bcftools concat -a -D*.

#### Somatic short indel calling with GATK4 Mutect2^30^

Somatic variants were called on prepared BAM files for the primary human chromosomes with *gatk* (v.4.1.5.0) *Mutect2*, where a ‘somatic variant’ is defined as one that exists in the tumor/experiment sample but not in the normal/control sample if in matched sample mode, or, in unmatched mode, as one not classified as a germline variant (considering allele frequency and other criteria). Importantly, *gatk* v4.1.5.0 was used rather than v.4.1.4.1, due to a bug in v4.1.4.1 that ends up inappropriately flagging reads as malformed, and breaking the run. Scattered VCFs were merged with *picard* (v.2.18.21) *MergeVcfs* and Mutect2 stats were merged with *gatk* (v.4.1.4.1) *MergeMutectStats*. Pileup summaries were ascertained for the RPE tumor analysis-ready BAM files with *gatk* (v.4.1.4.1) *GetPileupSummaries*, with GATK hg38 gnomad supplied as the population reference. Contamination table was calculated from the pileup summaries table with *gatk CalculateContamination*. Variants were scored with *FilterMutectCalls*, with Mutect2 stats and contamination table included in filtering. Variants that were not assigned a “PASS” were filtered out with *bcftools* (v.1.10.2) *view*. Interval-scattered VCFs were merged with *picard* (v.2.18.21) *MergeVcfs*. In-cohort samples were prepared for consolidation by consolidating their respective names with *bcftools reheader*. Finally, in-cohort sample VCFs were consolidated and de-duplicated with *bcftools concat -a -D*.

#### Variant call file finalization

For the RPE tumor sample, the germline and somatic VCFs were merged with *picard* (v.2.18.21) *MergeVcfs*.

### Genome individualization

#### Chromosome-wise patching of consensus reference genome with SNPs & indels from VCF

The GRCh38 human genome reference sequence was split up by chromosome with *samtools faidx* (v.1.10). Each chromosome was patched with SNPs and indels from the finalized VCF at specific loci using *bcftools consensus* (v.1.10.2). For the *consensus* command, the following parameters were invoked: -s <study_group> (tumor vs. normal or experiment vs. control), -H <haplotype> (haplotype={1, 2}; creates a bi-haplotype individualized genome), -p <haplotype> (prepends chromosome numbers with ‘{1, 2}_’ to keep track of haplotype), -c <output_chainfile> (to create a ‘chainfile’ that will be used for coordinate re-mapping later on). Finally, the full-length bi-haploid sequences are reconstituted from their chromosome-wise constituents via awk.

#### Coordinate lift-over of genome annotation

The genome coordinates of the *GENCODE v31* annotation GTF, originally corresponding to GRCh38, were transformed to the coordinate space of each of the two individualized haplotypes, using *CrossMap*^*31*^ *liftover* (v.0.3.9). (Please note that, as of now, this coordinate lift-over step causes a small amount of data loss in the resulting individualized annotation files; this occurs in the rare case where the genetic alterations for a given transcript are such that the program is unable to resolve an updated set of exon boundaries. However, as will be observed in *Results*, any effect this may have is evidently outweighed by the otherwise *improvement* in mapping performance as compared to when using an index generated from a reference genome such as GRCh38.) These annotation files (along with the individualized sequences) are subsequently used to generate genome indices for RNA mapping with *STAR*^*32*^.

### RNA sequencing analysis and transcriptome assembly

We used the workflow described in the original *ProteomeGenerator*^*17*^ paper, as summarized below.

#### Creation of individualized genome indices (1 per haplotype) for STAR

An index for the newly created custom genome was generated with *STAR* (v.2.7.3a) in *genomeGenerate* mode. Splice junction overhang threshold was given as (read length/2)-1 for paired-end reads, and (read length -1) for unpaired reads. Max genome suffix length was set to 1000. Other parameters were left at default.

#### RNA-seq mapping

RNA-seq input data (in FASTQ or BAM format) was mapped with *STAR*^*32*^ (v.2.7.3a), with read groups / biological replicates mapped separately. Two-pass mode was basic, chimeric reads were output within BAM, max intron and mate gap was set at 500,000 (for full list of user-specified parameters, see rule RNA_01_STAR_AlignRNAReadsByRG in ProteomeGenerator.py).

#### Transcriptome assembly

After a filtering step (removing low quality and multi-mapping (<=3) reads), remaining reads were assembled into transcript models via *StringTie*^*22*^ (StringTie, 37) (v.1.3.4). *StringTie* runs in either *de novo* (technically, genome-guided *de novo* to distinguish from genome-free assemblers like *Trinity*^*33*^) or annotation-guided mode. *De novo* mode was used in the present analysis. User-specified parameters including coverage threshold of 2.5, minimum transcript length of 210, minimum isoform fraction of 0.01. Following transcriptome assembly, all read group transcriptomes are non-redundantly merged (*StringTie merge*). *Merge* uses the same parameters as the assembly step, plus minimum transcripts per million (TPM) of 1 and inclusion of retained introns. These transcript models were subsequently read out to their corresponding cDNA sequences with *gffread*^*34*^ (0.9.12).

### Sample-specific proteome database constructions

For each cDNA sequence, 6-frame translation and open reading frame selection was performed with *TransDecoder*^*23*^ (v.2.5.0). In contrast with the original *ProteomeGenerator*, where only the single longest ORF was retained, PG2 retains all ORFs as specified by the user (default nucleotide length greater than 70*3), or that had significant BLAST homology (default expectation value > 1e-5; v.2.2.31). These default settings were implemented empirically after identifying greater numbers of peptides at equivalent global false discovery rate (FDR). Additionally, PG2 employed a customized version of *TransDecoder* that allows for adjustable overlap of retained ORFs. After translation and open reading frame selection, the selected candidate protein sequences were re-mapped onto their respective genomic coordinates using a script supplied by the *TransDecoder* distribution, and the re-mapped sequences are compiled to produce the PG2 sample-specific proteome FASTA.

### Protein mass spectrometry (MS/MS) search

Peptide fragmentation spectra were matched to the newly created PG2 database using MaxQuant^2^ (version 1.6.2.3) or PEAKS^1^ (version 8.0). MaxQuant was packaged into the release distribution of PG2, as it is Linux-compatible. The raw spectra files used in the present analysis were fractionated and digested with trypsin and/or Glu-C, as specified in the *Datasets* section. Parameters specified in the user configuration file (configuration files used in present analyses are supplied in the GitHub repository under configfiles/) are automatically propagated into the MaxQuant parameters XML file. The default template, mqpar_template.xml, is included in the GitHub repository. The template itself can be customized; if so, the custom template can be supplied to PG2 via input_files->proteomics_module->custom_params_xml in the PG2 configfile. PEAKS is included in the present analysis as its accuracy has been observed to be superior in recent studies.^17^

### Post-MS/MS analysis

A novel peptide mapping and protein expression analysis suite was developed for the present analysis and included in the release distribution. For the novel peptide mapping functionality, identified peptides were cross-referenced against the reference proteome to identify non-canonical peptides. With the assistance of the *snpEff*^*35*^ variant annotation tool, novel peptides were then remapped to the genes and transcripts, and ultimately the mutations from which they derive, which were compiled and tabulated in a user-friendly manner (*novel_analysis* directory). For initial protein expression analysis, a script was written that compares, between matched experimental and control conditions, the number of mass spectrometry-identified peptides for each gene (or de novo isoform).

### Reagents

Mass spectrometry grade (Optima liquid chromatography–mass spectrometry, LC–MS) water, acetonitrile (ACN), and methanol were purchased from Fisher Scientific (Fair Lawn, NJ). Formic acid of >99% purity (FA) was obtained from Thermo Scientific. All other reagents at MS-grade purity were obtained from Sigma-Aldrich (Saint Louis, MO).

### Cell lines

Human AML cell lines, Kasumi1, MV411, U937, OCI-AML2 and OCI-AML3, were obtained from the American Type Culture Collection (ATCC, Manassas, Virginia, USA) or DSMZ (Leibniz Institute, Braunschweig, Germany). These cell lines were cultured in RPMI-1640 medium (Corning) supplemented with 10% fetal bovine serum (Gibco), 100 U/ml penicillin and 100 μg/ml streptomycin in a humidified atmosphere at 37 °C and 5% CO_2_. All cell lines were authenticated by STR genotyping (Integrated Genomics Operation, Center for Molecular Oncology, MSKCC). The absence of Mycoplasma species contamination was verified using MycoAlert Mycoplasma detection kit, according to manufacturer’s instruction (Lonza). Human K052 cells (first described and utilized in Cifani et al., 2018) were obtained from the Japanese Collection of Research Bioresources Cell Bank, identity confirmed using STR genotyping (Genetica DNA Laboratories, Burlington, NC), and cultured as described.^17^ Cells were collected while in the exponential growth phase, washed twice in ice-cold phosphate-buffered saline, snap-frozen, and stored at -80 °C.

### Decitabine treatment

5-Aza-2’-deoxycytidine (Decitabine, Sigma A3656) was dissolved in DMSO at 100 mM. Cells were treated with 300 nM Decitabine or DMSO control for 2 days. After another 5-day culture, the cells were collected for protein and RNA extraction.

### RNA sequencing (sample preparation and sequencing)

RNA (from the 5 grouped AML cell lines) at 5 million cells per sample was extracted using the RNeasy mini kit (QIAGEN) according to the manufacturer’s instructions. After RiboGreen quantification and quality control by Agilent BioAnalyzer (Agilent, Santa Clara, CA, USA), 500 ng of total RNA with RIN values of ≥10 underwent polyA selection and TruSeq library preparation according to the manufacturer’s instructions (Illumina TruSeq Stranded mRNA LT Kit, RS-122-2102), with 8 cycles of PCR. Samples were barcoded and sequenced using HiSeq 4000 instrument in a pair-end 100-bp run, using the HiSeq 3000/4000 SBS Kit (Illumina). An average of 121 million paired reads was generated per sample. Ribosomal reads represented 0.9-2.8% of the total reads generated and the percent of mRNA bases averaged 68%. RNA-seq data for the K052 cell line, was obtained as previously described.^17^

### Whole genome sequencing (sample preparation and sequencing)

Genomic DNA from 5 million cells per sample was extracted using the PureLink Genomic DNA mini kit (Invitrogen) according to the manufacturer’s instructions. After PicoGreen quantification and quality control by Agilent TapeStation (Agilent, Santa Clara, CA, USA), 500 ng of genomic DNA were sheared using a LE220-plus Focused-ultrasonicator (Covaris catalog # 500569) and sequencing libraries were prepared using the KAPA Hyper Prep Kit (Kapa Biosystems KK8504) with the following modifications. Libraries were subjected to a 0.5X size selection using AMPure XP beads (Beckman Coulter, A63882) after post-ligation cleanup. Libraries were not amplified by PCR and were pooled at equal volume and quantitated based on their initial sequencing performance. Samples were sequenced using NovaSeq 6000 instrument in a pair-end 150-bp run, using the NovaSeq 6000 SBS v1 Kit and an S2 flow cell (Illumina) with target mean 40x coverage.

### Protein extraction and proteolysis

Fifty million cells per sample were collected and washed with cold PBS. Cell pellets were resuspended in 6M guanidinium hydrochloride (GdnHCl) and 100 mM ammonium bicarbonate (ABC), and lysed using the E210 adaptive focused sonicator (Covaris, Woburn, CA) for 10 min at 4 °C. Extracted proteins were reduced with 10mM dithiothreitol (DTT) and alkylated with 55mM iodoacetamide (IAA), and IAA was quenched with DTT. The protein content in cell lysate was determined using the BCA assay according to the manufacturer’s instructions (Pierce, Rockford, IL). One mg of each sample was applied for trypsin proteolysis. Samples were diluted in 100 mM ammonium bicarbonate and digested with LysC endopeptidase (Wako Chemical, Richmond, VA) at 1:100 w/w (protease:proteome) at 37 °C for 6 hours and MS sequencing-grade modified trypsin (Promega, Madison WI) at 1:50 w/w (protease:proteome) at 37 °C overnight, followed by 1:33 w/w trypsin digestion for additional 4 hours. Another 1 mg of each sample was applied for Glu-C proteolysis. Samples were diluted in 100 mM ammonium bicarbonate and digested with MS sequencing-grade Glu-C (Promega, Madison WI) at 1:50 w/w (protease:proteome) at 37 °C overnight, followed by 1:33 w/w Glu-C digestion for additional 6 hours. Proteolysis was quenched with 1% trifluoroacetic acid (Sigma-Aldrich, Saint Louis, MO), and digested peptides were purified using solid-phase extraction (SPE) using C18 Macro Spin columns (Nest Group, Southborough, MA). SPE eluates were lyophilized and stored at −20 °C until analysis.

### Chromatographic peptide fractionation

Purified peptides were resuspended in 200 μL of 0.1% formic acid in water (0.5 μg/μL) and injected onto the strong cation exchange (SCX) column (Protein Pak Hi-Res SP 7 μm 4.6 × 100 mm, Waters) at a constant flow rate of 250 μL/min with a column temperature of 25 °C using the Alliance high-performance liquid chromatography (HPLC) system (Waters, Milford MA). Gradient was run with 5% acetonitrile and 0.1% formic acid in water for solvent A and 1 M potassium chloride in 5% acetonitrile and 0.1% formic acid in water for solvent B. Column gradient was set as follows: 0–3 min (0% B), 3–83 min (0–35% B), 83–90 min (35-50% B), 90-91 min (50-100% B), 91-100 min (100–0% B), and 100–120 min (0% B). In total, six fractions were collected from 3 to 75 min per sample for each condition. Fractions were then purified using solid phase extraction (SPE) with C18 Macro spin columns according to manufacturer’s instructions (Nest Group). SPE eluates were lyophilized and stored at −20 °C until analysis.

### Nanoscale liquid chromatography and nanoelectrospray Orbitrap mass spectrometry

Nanoscale liquid chromatography was performed using the Dionex UltiMate 3000 HPLC system equipped with NCS-3500RS nano/capillary pumps (Dionex). Each fractionated sample was resuspended in 20 μL of 0.1% formic acid in water, and 2 μL was loaded on an Acclaim PepMap 100 C18 trap column (100 μm x 20 mm, Thermo Scientific) at 3 µL/min. Separation was achieved using EASY-Spray HPLC Columns (75 μm x 500 mm, 2 μm ultrapure silica, Thermo Scientific). Solvent A was 0.1% formic acid in water, and solvent B was composed of 0.1% formic acid and 80 % acetonitrile in water. Column gradient was set as follows: 0-7 min (2% solvent B), 7-80 min (2-33% solvent B), 80-157 min (33-40% solvent B), 157-158 min (40-90% solvent B), 158-170 min (90% solvent B), 170-170.2 min (90-1% solvent B), and 170.2-185 min (1% solvent B). The Q-Exactive HF mass spectrometer (Thermo Scientific) was used for peptide analysis. Precursor ions in the 400-1500 m/z scan range were isolated using the Quadrupole and recorded every 3 seconds using the Orbitrap at 60,000 resolution with an automatic gain control (AGC) target set at 3 x 10^6^ ions and a maximum injection time of 54 ms. Top20 data-dependent MS2 precursor ion selection was enforced, limiting fragmentation to monoisotopic ions with charge 2–4 and precursor ion intensity greater than 1.6 x 10^5^, and dynamically excluding already fragmented precursors for 15 seconds (10 ppm tolerance). Selected precursor ions were isolated (Q1 isolation window 0.7 m/z) and fragmented using Higher-energy C-trap dissociation (HCD, normalized collision energy 30) using the top speed algorithm. Product ion spectra in the 200-2000 m/z scan range were recorded in the Orbitrap at 30,000 resolution (AGC of 2 × 10^5^ ions, maximum injection time of 50).

### Databases

The consensus human genome reference GRCh38.p11^36^, GENCODE^37^ version 31 genome annotation and translated transcriptome annotation, and consensus human proteome reference UniProt^38^ (SwissProt version 9606, concatenation of canonical sequences + isoforms + cRAP^39^ contaminants database) were used in the experiments.

### Artificial introduction of SNPs to test PG2

The original bam files were mutated using a python script whereby the allele frequencies of the mutations were set to either 1.0 or 0.5 for homozygous and heterozygous mutations respectively. Successful introduction of mutations was confirmed manually by using IGV. The resulting mutated bam files were used as input bam files. The detection of mutation was checked in the output vcf files and the final proteome output file.

### Automated validation framework

Briefly, the same VCF used to create the individualized genome was also passed to SnpEff^35^, along with the genome and translated transcriptome annotation (GENCODE^37^ v31). After building a database out of the supplied information SnpEff parses all variants in the VCF, generating an annotated VCF containing information about each known transcript that a given variant is expected to impact. For each annotated transcript’s proteoform sequence, the variant context was extracted; that is, a subset of the full sequence centered around the position of the variant. This was done in order to maximize the probability that the ensuing comparison would differ only with respect to the variant in question (complete transcript sequences are likely to have multiple alterations compared to their canonical counterparts). Next, this variant context was compared by local alignment (via Biopython^40^ pairwise2) to proteoforms in PG2’s sample-specific database that have been identified as bearing strong homology (e-value < 1e-40 via BLASTp^41^) to the annotated transcript from which the variant context in question derives. If the local alignment contains the exact anticipated amino acid change, then the variant is considered to be successfully recovered. Often, the homologous PG2 proteoform would not contain the variant context in question, and in these cases the resulting local alignment would be nonsensical. The absence of variant context within an ostensibly homologous proteoform may be due a skipped or truncated exon (which indels often cause directly) or simply an incorrect annotation-to-PG2 proteoform mapping by BLASTp. Missing variant contexts were not useful for validating variant propagation and are therefore excluded from analysis. As such, ‘failed’ recoveries were those for which the relevant variant context *was* inspectable in both haplotypes (i.e. those whose variant context is appropriately recovered by local alignment via 1 or more associated proteoforms per haplotype), but the predicted amino acid change occurs in neither. All recoveries marked as failures by automated validation are then inspected manually.

### Code and data availability

All code is openly available via https://github.com/kentsisresearchgroup/ProteomeGenerator2, which includes default YAML configuration files and instructions for PG2 installation and use. Mass spectrometry data have been deposited to the ProteomeXchange Consortium via the PRIDE partner repository with the data set identifier PXD037061. Whole genome sequencing and RNA sequencing data have been deposited in the NCBI Sequence Read Archive with the BioProject accession number PRJNA878804. Processed files are available from Zenodo (10.5281/zenodo.7068642). Mass spectrometry raw data and RNA sequencing data for K052 were obtained from previously published data sets PXD008381 and PRJNA427716, respectively^17^.

## Results

### Validation of the sample-specific bi-haploid genome assembly

PG2 builds on the ProteomeGenerator^17^ framework to integrate genome DNA sequencing and variant calling with referenced or de novo transcriptome assembly to enable the identification of genetic and mutational variants and non-canonical protein isoforms (**Figure 1**). This approach requires sample-specific genome customization. As little has been reported to-date on the use of *bcftools* consensus for genome customization, we first sought to confirm that the sample-specific bi-haplotype genomes deployed in PG2 constitute valid genome assemblies, comparable at a minimum to the reference assemblies from which they derive. To that end, each haplotype of the individualized genome for the Kasumi1 cell population (KASUMI_H1 and KASUMI_H2) was benchmarked against GRCh38 using the QC tool Quast^42^. KASUMI1_H1 and KASUMI1_H2 demonstrated genome fractions of 99.993% and 99.995% respectively, where genome fraction was defined as the proportion of bases in the reference assembly for which there are aligned bases in the test assembly. GC content was 41.00% and 40.99% respectively, as compared to 40.99% in GRCh38. There were 7 misassembled contigs for a total length of 9842124 bases in KASUMI1_H1 and 9 misassembled contigs for a total 14771393 bases in KASUMI1_H2. There were no fully unaligned contigs in either haplotype; there were 3 partially unaligned contigs corresponding to 5264 unaligned bases in KASUMI1_H1, and 4 partially unaligned contigs corresponding to 9713 unaligned bases in KASUMI1_H2. In comparison to GRCh38, there were 1764450 total mismatches and 436238 total indels in KASUMI1_H1, and 3680456 total mismatches and 793844 total indels in KASUMI1_H2. The rate of N’s (i.e., uncalled bases) per 100 kilobases in the two test assemblies were 4985 and 4985 respectively, versus 5324 in GRCh38. Out of 2881886 genomic features (i.e., genes, transcripts, and CDS) included in the GRCh38 GENCODE v31 annotation, 2879325 (99.91%) and 2878590 (99.88%) were completely aligned in each respective test assembly, and another 687 (0.02%) and 1417 (0.05%) were partially aligned (i.e., at least 100bp of feature were aligned). Complete QUAST metrics for all datasets are included in the supplementary data. Thus, PG2 enables sample-specific bi-haploid genome assembly, which is expected to improve with the implementation of improved sequence alignment algorithms and genome assemblies.

**Figure 1.**
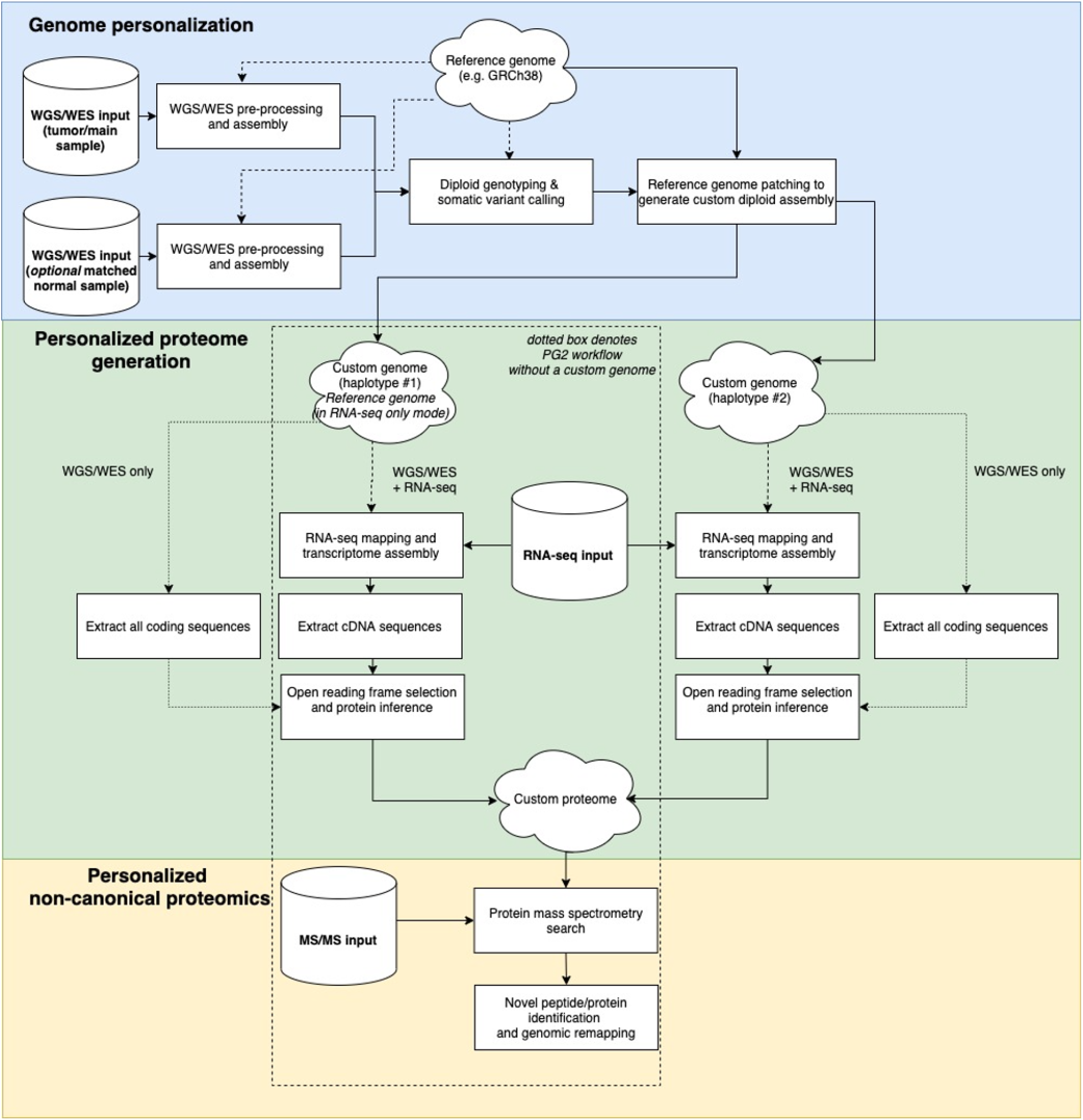
ProteomeGenerator2 workflow. PG2 maps DNA sequencing reads to reference genomes, identifies germline and somatic variants using flexible variant callers, and then modifies the reference genome to incorporate called variants, yielding an individualized or specimen-specific genome assembly. Subsequently, PG2 maps RNA sequencing reads to the custom genome, assembles the transcriptome as the user specifies (*de novo* hybrid or genome-guided), and assembles resultant open reading frames into a comprehensive, personalized protein sequence database (the PG2 proteome). This database becomes the search space for specimen-specific mass spectrometry analysis. Following mass spectrometric analysis, non-canonical peptides are aggregated and If no DNA sequencing data is available, PG2 can be used in RNA-only mode, utilizing the reference genome (*a la* PG1, as is encapsulated in the dotted box).

### Impact on RNA-seq mapping and transcriptome assembly

The sample-specific genome assembly is intended to be a more accurate (given its incorporation of sample-specific variants) and complete (given that its sequential format better approximates non-haploid genotypes) representation of the sample organism, as compared to the consensus reference assembly. As such, we reasoned that the use of the sample-specific assembly could produce improvements in downstream genome-referenced analyses, and in the case of PG2, RNA read mapping and transcriptome assembly.

To that end, we first examined the impact of genome individualization on RNA-seq mapping using *STAR*^*32*^ (see Methods for parameters used), comparing the mapping accuracy of the customized genomes to that of the *GRCh38* reference genome. Bi-haplotype genomes were generated for each of the 5 different human acute myeloid leukemia (AML) cell lines, and corresponding raw RNAseq reads for the respective lines were mapped to each of the two sample-specific haplotypes as well as the GRCh38 genome (**Figure 2**). Uniformly across all 5 cell lines, both sample-specific haplotypes had higher proportion of uniquely mapped reads in comparison to GRCh38 (88.02% for *AML_H1* vs 87.82% for GRCh38, *p* = 0.007; 88.20% for *AML_H2* vs 87.82% for GRCh38, *p* = 0.0003; **Figure 2a-b**) and longer mean lengths of mapped reads (198.96 for *AML_H1* vs 198.68 for GRCh38, *p* = 0.0001; 198.96 for *AML_H2* vs 198.68 for GRCh38, *p* = 0.0001; **Figure 2d**) as well as lower proportions of multi-mapped reads (6.89% for *AML_H1* vs 7.01% for *AML_H2, p* = 0.072; 6.73% for *AML_H2* vs 7.01% for GRCh38, p=0.0016), chimeric reads (2.27% for *AML_H1* vs 2.29% for GRCh38, *p* = 0.0039; 2.27% for *AML_H2* vs 2.29% for GRCh38, *p* = 0.0039) and unmapped reads (4.47% for *AML_H1* vs 4.53% for GRCh38, p=0.004; 4.47% for *AML_H2* vs 4.53% for GRCh38, *p* = 0.004; **Figure 2c**). Although these findings are reported here in terms of their aggregate mean values across cell lines, it should be noted that the association held uniformly in every individual comparison within each cell line.

**Figure 2.**
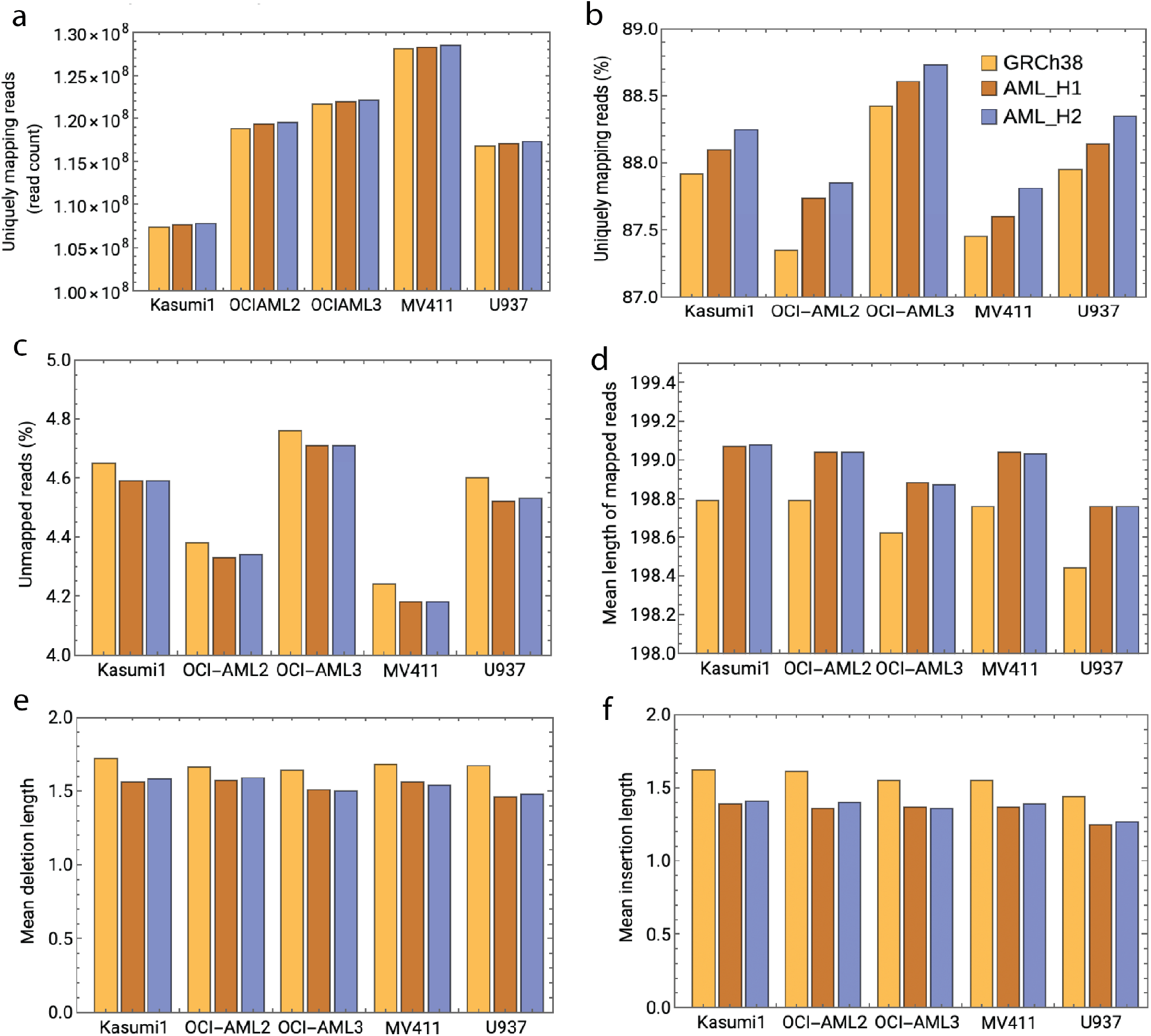
Comparison of RNA read mapping metrics with PG2-generated custom haplotypes versus GRCh38. Custom bi-haplotype genomes for each of the 5 AML cell lines (denoted AML_H1 and AML_H2) were generated and used for RNAseq read mapping with STAR aligner. With respect to mapping success rate, both custom haplotypes achieved (versus GRCh38) higher percentages of uniquely mapped reads, lower percentages of unmapped reads, and higher average length of mapped reads, in every comparison. With respect to mapping error rate, both custom haplotypes achieved lower mismatch rates, lower average length of insertion, and lower average length of deletion, again in every comparison.

To evaluate per-base mapping performance, we examined the mismatch rate per base, average length of insertion (referring to when an RNA-seq read exhibits inserted bases in comparison to its mapped location on the reference genome) and average length of deletion. With respect to each of the error rate metrics, both *AML_H1* and *AML_H2* produced significantly lower error rates on average than did *GRCh38*, which was consistent within every individual comparison within each cell line (**Figures 2e-f**). Taken together, these findings collectively support the notion that genome customization improves RNA read mapping accuracy.

### Proteogenomic propagation and recovery of known genetic variants

Having verified the soundness of the sample-specific genome assembly approach as well as its improvements for integrative analyses, we next sought to examine PG2’s underlying end-to-end functionality of propagating genomic variants to sample-specific predicted proteome sequences. To this end, we analyzed targeted DNA sequencing (MSK-HemePACT targeted DNA assay^43^) and bulk RNA sequencing data obtained from human leukemia K052 cells that are known to harbor a heterozygous genetic mutation in exon 1 of *SRSF2* (corresponding to P to H missense substitution at codon position 95 of the canonical proteoform).^17^ Assuming correct functionality, it would be expected that the heterozygous mutant allele would be disseminated to one of the two individualized haplotypes and therefore, given the requisite mRNA expression of *SRSF2* in this sample for de novo assembly of the transcript, we would expect both the wildtype (P) and mutant (H) translated amino acid sequence isoforms of SRSF2 to appear in the proteome. Indeed, this was observed (all output proteomes and pertinent processed files are available on Zenodo).

### Proteogenomic propagation and recovery of synthetically introduced variants

To test the performance of PG2 in detecting single nucleotide variants, we introduced synthetic variants into the DNA sequencing data of the K052 cell line within the region that encodes *BRCA1* using jvarkit^44^. We introduced mutations in amino acid coding regions, in non-coding regions, in mutations that result in no amino acid change (silent mutations), as well as one mutation affecting a stop codon and one mutation affecting a start codon (**Figure 3a**). While introducing the mutations in the bam file of the K052 cell line, we used an allele frequency of 0.5 to simulate heterozygous mutations as well as an allele frequency of 1 to simulate homozygous mutations. The seven missense mutations in the coding regions were then propagated to the final proteome fasta file using PG2. The unmodified (or “wildtype”) K052 dataset computation resulted in 4 isoforms of BRCA1, the homozygous manipulated bam file resulted in 7 isoforms of BRCA1, each of them containing an isoform with an artificially introduced mutation and none of the wildtype BRCA1 isoforms. The heterozygous K052 dataset computation resulted in 11 isoforms (representing the wildtype isoforms plus the mutant isoforms; **Figure 3b**). Appropriately, mutant start and stop codons were found in the generated vcf file, and were filtered out after the mutect2 step of the PG2 pipeline. Similarly, non-coding and synonymous substitutions produced no changes in the predicted proteomes. These results indicate that PG2 is able to accurately detect mutations and integrate them in the sample-specific predicted proteomes.

**Figure 3.**
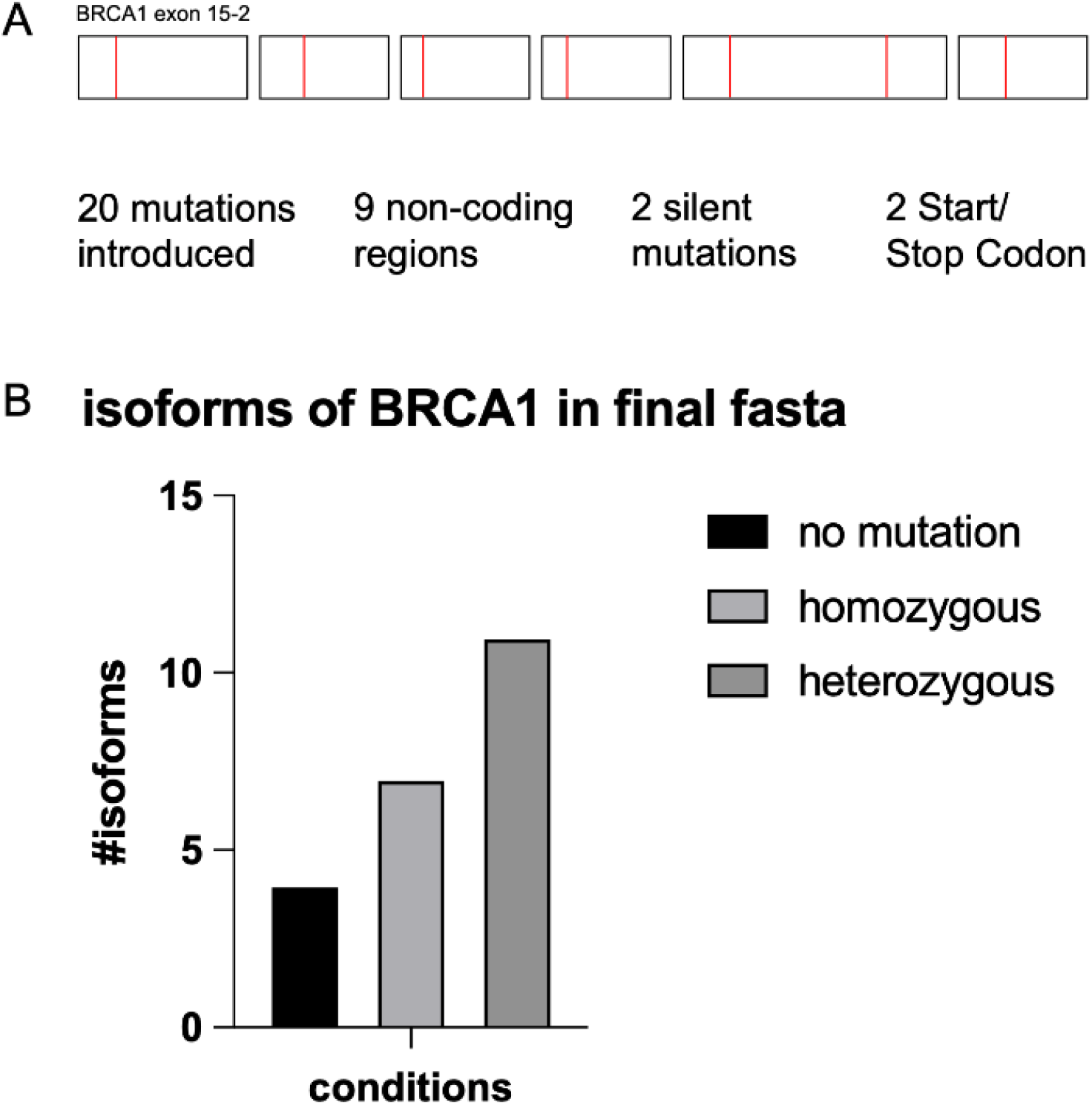
Detection of artificially introduced SNPs in the BRCA1 gene locus of the K052 cell line. Seven missense mutations in amino acid coding regions that are not a stop or start coding were introduced. All of them are found as amino acid changes in the final fasta file. Allele frequencies of mutations were set to 0.5 (heterozygous) or 1 (homozygous). Without any mutation, 4 isoforms of BRCA1 are in the final fasta file. In the condition with the mutations introduced with a heterozygous AF, 11 isoforms are in the final fasta file. In the fasta file with the homozygously introduced mutations, 7 isoforms of BRCA1 are found in the final proteome (only the one containing mutations, not the original “WT run” isoforms. SNPs that are in STOP or START codon are found in the vcf file however, filtered out after the mutect2 step. SNPs that are in noncoding regions are also filtered out. Silent mutations are found in the vcf file also after the filtering step however, do not result in a new protein isoform.

### Genome-wide validation of end-to-end genetic variant proteogenomic analysis in human AML cell lines

Encouraged by the results of manual validation, we next developed an automated validation framework to examine variant-containing PG2 proteome sequences at genome scale. This was accomplished by annotating the genomic VCF files with variant amino acid effect predictions made by SnpEff, and subsequently cross-referencing these predictions against proteoform sequences assembled orthogonally by PG2 (Supplemental Figure 1). In addition to serving as method validation, this particular module compiles identified variants for PG2 users, at both the transcript and protein levels. These variant results are indexed by gene, annotated transcript with positional change and associated PG2-assembled proteoforms.

Using this approach, we performed integrative proteogenomic analysis of five human AML cell lines using matched whole-genome DNA sequencing, transcriptome RNA sequencing, and multi-dimensional separation and multi-protease digestion high-resolution mass spectrometry proteomics (see Methods). We used SnpEff to annotate variants that are expected to result in missense substitutions, in-frame insertions, in-frame deletions, or frameshift sequences. We required that at least one transcript, in which the variant is known to reside, be assembled in both haplotype calculations of PG2, as governed by *de novo* transcriptome assembly (**Figure 1**). In this way, we could ensure that a failure to recover a given (e.g., heterozygous) variant was due to a technical failure of PG2 and not, for instance, due to the variant residing in the unassembled (and thus, non-inspectable) transcript of the opposite haplotype. Additionally, candidate variants denoted by the automated framework as failed recovery events were manually inspected for further confirmation, which allowed for ‘rescue’ retrieval of some variants, as well as designation of some variants as non-inspectable/unrecoverable from a validation standpoint (due, for instance, to the variant-containing portion of a transcript being absent from the assembled protein sequence, or it residing in a highly repetitive region where inspection is not possible). Importantly, manual inspection is only needed because of technical limitations of the automated validation method, and does not impact the underlying data. The full specification of the validation method is described in the Methods section.

Using automated validation, we found that PG2 successfully recovered 3880/3907 (99.3%), 3548/3574 (99.2%), 3529/3556 (99.2%), 3627/3648 (99.4%), and 3503/3520 (99.5%) variants in the Kasumi1, OCI-AML2, OCI-AML3, MV411, and U937 cell lines, respectively (**Figure 4**). Inclusive of manual confirmation, 3905/3907 (99.9%), 3570/3573 (99.9%), 3551/3554 (99.9%), 3645/3647 (99.9%), 3518/3518 (100.0%) were successfully recovered. With respect to in-frame insertions, PG2 successfully recovered 53/55 (96.3%), 51/55 (92.7%), 62/65 (95.3%), 66/70 (94.2%), and 58/61 (95.1%) variants in the Kasumi1, OCI-AML2, OCI-AML3, MV411, and U937 cell lines, respectively. Inclusive of manual confirmation, 53/53 (100.0%), 54/55 (98.1%), 65/65 (100.0%), 66/68 (97.1%), and 58/60 (96.7%) were successfully recovered. With respect to in-frame deletions, PG2 successfully recovered 70/73 (95.8%), 79/80 (98.8%), 77/81 (95.1%), 80/81 (98.8%), and 68/72 (94.4%) variants in the Kasumi1, OCI-AML2, OCI-AML3, MV411, and U937 cell lines, respectively. Inclusive of manual confirmation, 70/70 (100.0%), 79/79 (100.0%), 77/78 (98.7%), 80/80 (100.0%), and 69/69 (100.0%) were successfully recovered. With respect to frameshift events, PG2 successfully recovered 109/119 (91.6%), 92/95 (96.8%), 97/106 (91.5%), 82/86 (95.3%), and 90/98 (91.8%) variants in the KASUMI1, OCI-AML2, OCI-AML3, MV411, and U937 cell lines, respectively. Inclusive of manual confirmation, 114/117 (97.4%), 91/93 (97.8%), 100/103 (97.1%), 84/85 (98.8%), and 95/96 (99.0%) were successfully recovered. Thus, PG2 enables genome-wide integrative proteogenomics.

**Figure 4.**
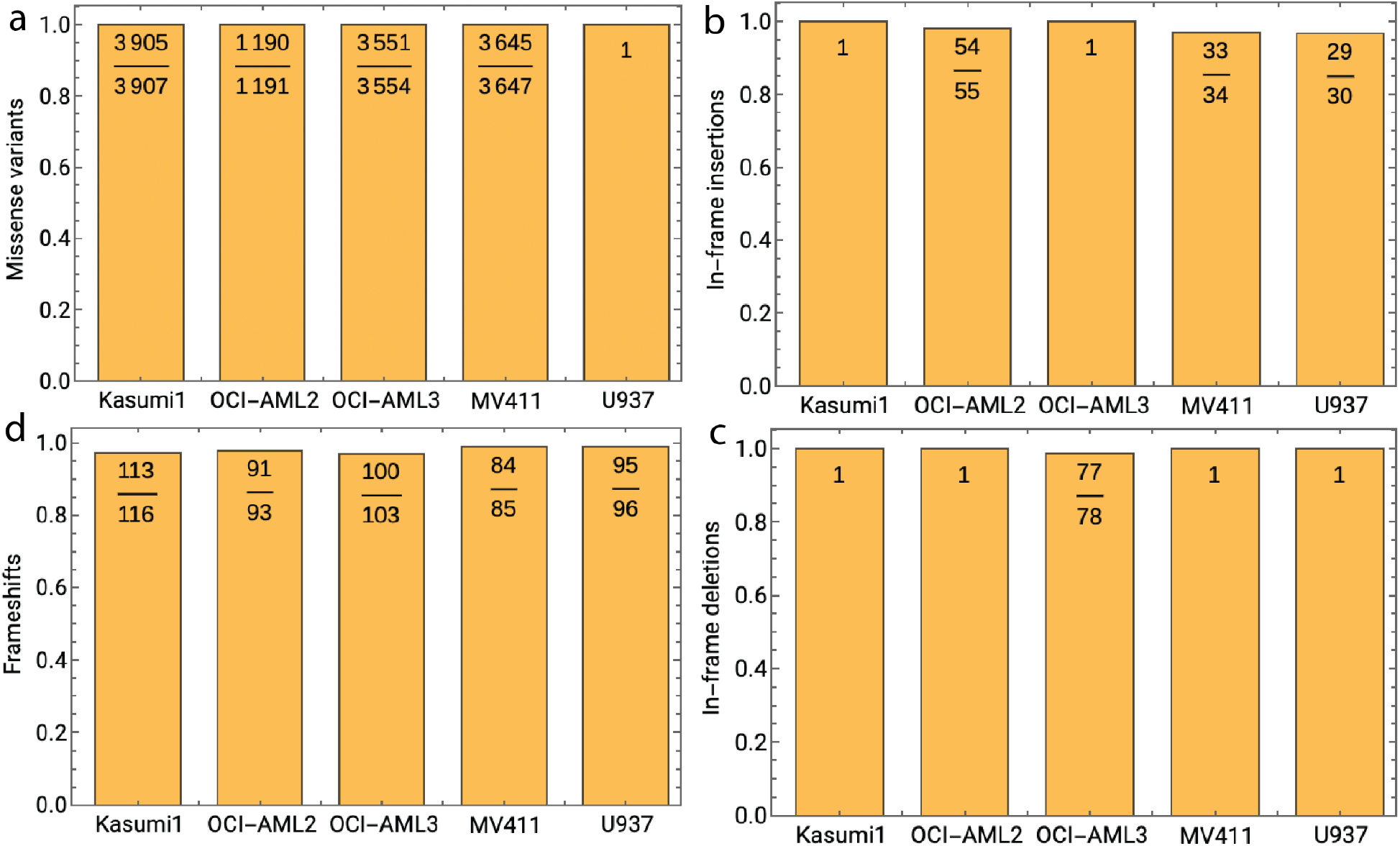
Genome-wide end-to-end validation of propagation of DNA sequencing mutations, from variant call file (VCF) to specimen-specific protein sequence database (FASTA). Variant call files for each of the 5 AML cell lines were annotated using *SnpEff* to generate predictions of impact on amino acid sequence. Subsequently, these predictions were validated on the proteome side at genome-scale by automated validation, supplemented by manual inspection. The depicted bar graphs illustrate (clockwise from the top left) variant propagation success rates of a) missense substitutions, b) in-frame insertions, c) in-frame deletions, and d) frameshift mutations. The complete validation framework is available in the Supplemental Figure 1.

### Identification of variant protein isoforms using integrative proteogenomics

We reasoned the integrative proteogenomic framework of PG2 would enable the identification of variant protein isoforms. For example, CCAAT/enhancer-binding protein alpha (CEBPA) is known to encode at least two protein isoforms, p42 and p30, which differ in that p30 begins with an internal start codon downstream of p42’s translation initiation site; p30 is therefore a subsequence of p42.^45^ The relative abundance of p42 and p30 has significant ramifications with respect to hematopoietic cell development; indeed, a significant fraction of acute myeloid leukemia patients have CEBPA mutations, with frameshift or nonsense mutations upstream of the p30 initiation site (codon 119 of p42) being an especially common alteration.^46,47^ The K052 leukemia cells used in the present study are known to harbor one such (heterozygous) nonsense mutation at codon 50 of the p42 proteoform. In order to correctly identify and document both isoforms – a C-terminally truncated p42 and a wildtype p30 – the assembler needs to be cognizant of the stop gained variant (otherwise outputting a full-length p42) as well as capable of generating multiple open reading frames per protein (otherwise typically generating only the single longest sequence, which in this mutant case is now p30). PG2 analysis revealed wildtype p42 isoform (occupying one haplotype), the C-truncated p42 (on the other haplotype), as well as wildtype p30 (**Figure 6a**). Thus, PG2 successfully addressed both issues, accounting for variants as well as offering support for multi-reading frames as a user-specified parameter (enabled by default).

**Figure 5.**
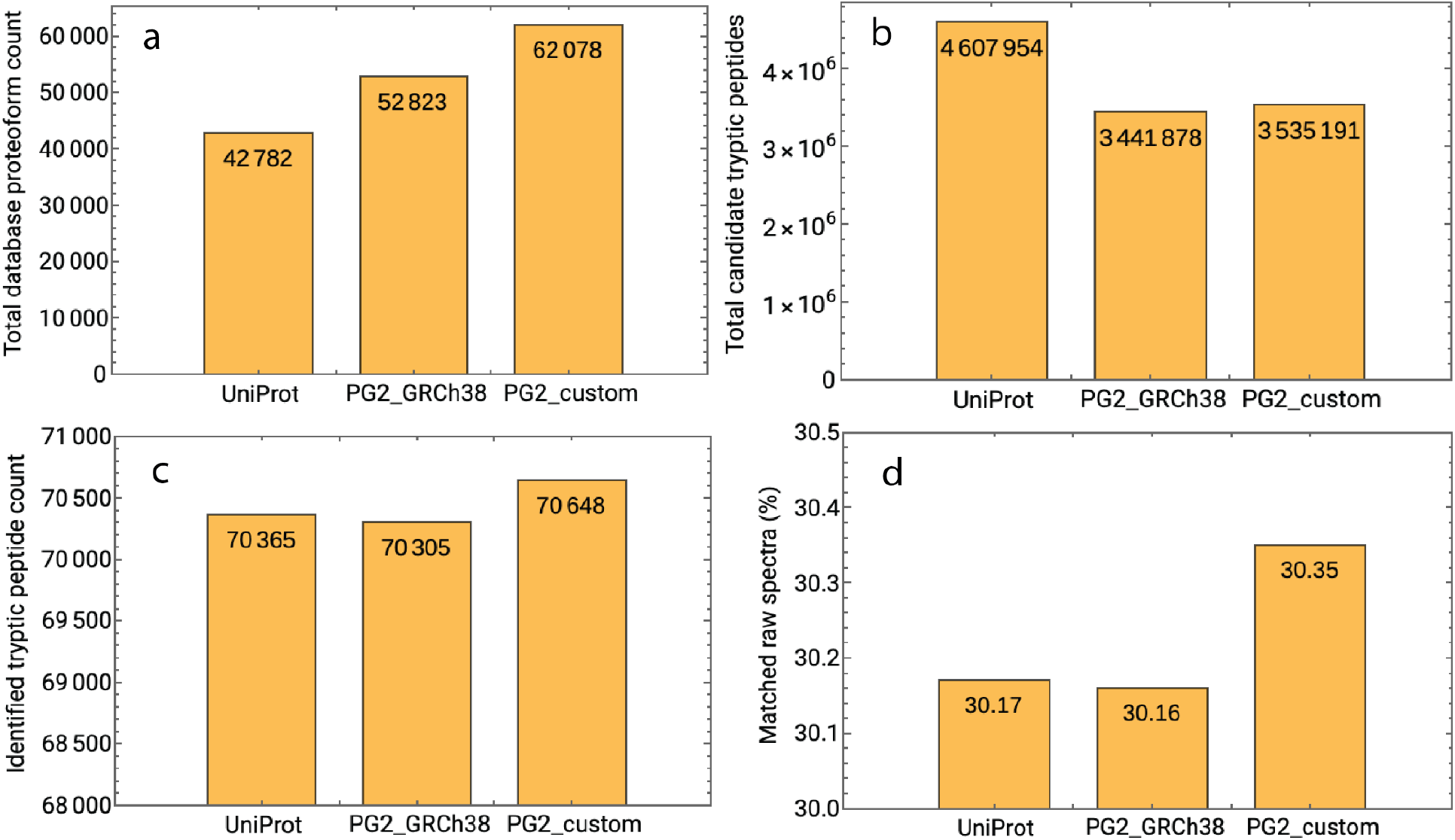
Proteomics performance of the PG2 proteome databases. Utilizing the KASUMI1 cell line proteogenomic dataset, two PG2-generated proteomes - one referencing an individualized genome (PG2_custom) and one referencing the reference genome GRCh38 (PG2_GRCh38) - were evaluated on the basis of database content and mass spectrometry performance, with the UniProt reference proteome as a benchmark. a) Total count of protein isoform entries in the proteome database. b) Total *in-silico*-calculated candidate enzyme-digested peptides (trypsin + Glu-C) for each respective database. c) Count of unique enzyme-digested peptides identified by mass spectrometric analysis. d) Percentage of successfully matched raw mass spectra.

**Figure 6:**
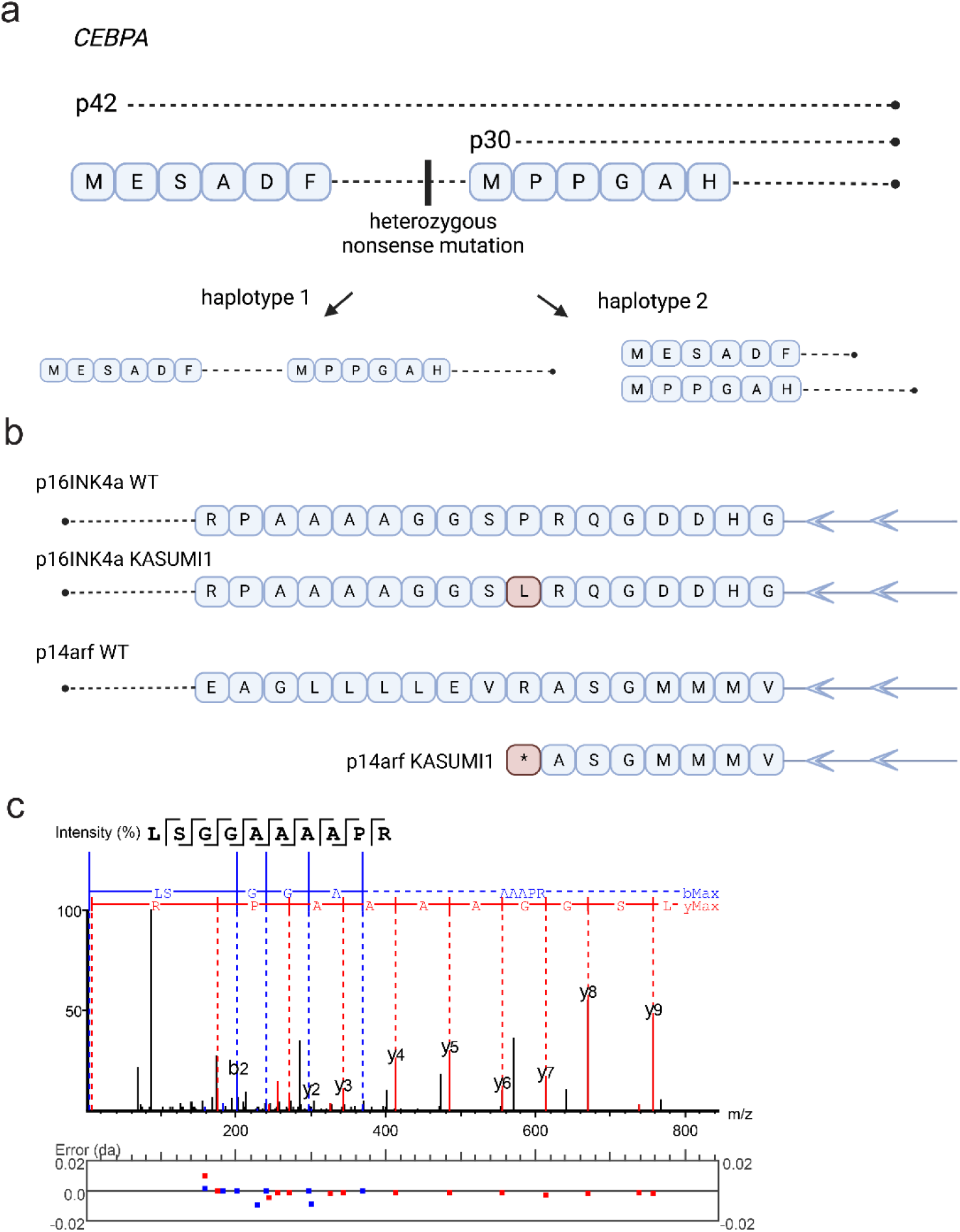
PG2 enables precise proteomic analysis of complex genotypes. a) The CEBPA gene encodes two distinct proteoforms, p42 and p30, which must be independently accounted for and aggregated, as it is the expression ratio between the two that is thought to drive oncogenesis. The K052 cell line from the present study harbors a heterozygous premature stop codon in p42, upstream of the p30 start codon. By employing multiple haplotypes, being variant aware, and allowing for outputting of multiple open reading frames per transcript, PG2 capably accounts for this complex genotype, as illustrated. b) The two protein-coding reading frames of the tumor suppressor gene CDKN2A: p16ink4A (p16) and p14arf (p14). A single nucleotide substitution induces a missense substitution in p16 and a nonsense codon in p14. c) p16 is identified uniquely by the non-canonical peptide LSGGAAAAPR, containing the P->L missense change.

Another striking example is that of the tumor suppressor gene CDKN2A, which actually encodes two different tumor suppressor proteins, *p16* and *p14*, whose overlapping reading frames are simply offset by 1 nucleotide. Interestingly, the Kasumi1 AML cell line harbors a SNP in CDKN2A that simultaneously encodes a missense mutation in p16 and a nonsense mutation in p14 (**Figure 6b**). Using PG2, we detected a p16-derived peptide, which contains the substituted amino acid, but no peptides associated with p14 (**Figure 6c**). This provides information that could suggest that, for example, the activity of mutated p16 is preserved whereas p14 is lost in certain forms of AML. Once again, the individualized and multi-reading frame-supporting approach of PG2 allows for such nuanced observations to be made.

### Non-canonical proteome investigation and differential comparison

The underlying goal of PG2 is to be able to discover the complete non-canonical proteome, spotlighting previously unrecognized targets for further study in an unbiased manner. To explore this utility, we analyzed matched whole-genome DNA sequencing, RNA sequencing, and mass spectrometry proteomics data obtained from Kasumi1 AML cells before and after treatment with the hypomethylating agent decitabine (DAC), currently used for treatment of AML.^48^ Using PG2’s matched experiment/control mode to perform harmonized calculations of the two conditions with a unified sample-specific genome, we generated in parallel sample-specific PG2 proteomes for the respective conditions and subsequently used them as reference sequences for mass spectral matching using the PEAKS algorithm (see Methods). Consistent with the expected activity of DAC to alter gene expression, we observed significantly increased global protein expression, with 3708 and 3222 proteins having increased and decreased mean peptide spectral intensities in the DAC-treated as compared to the DMSO treated cells, as normalized to housekeeping proteins ACTB, GAPDH, VCL, and LMNB1, respectively (**Supplemental Figure 2**).

Across both conditions, there were 374 proteins that were detected by at least one non-canonical peptide (i.e., one without an exact sequence match in the UniProt database), and 8 proteins that were identified exclusively by non-canonical peptides (**Figure 7a**). For example, we identified non-canonical peptides for CATSPERZ (**Figure 7b an 7c**), the abundance of which was modulated by DAC treatment. While the detailed extent and functional significance of acquired alterations in AML protein expression and its modulation by DAC are beyond the scope of current work, these results demonstrate the suitability of PG2 for nuanced investigation of the non-canonical or sample-specific proteome.

**Figure 7.**
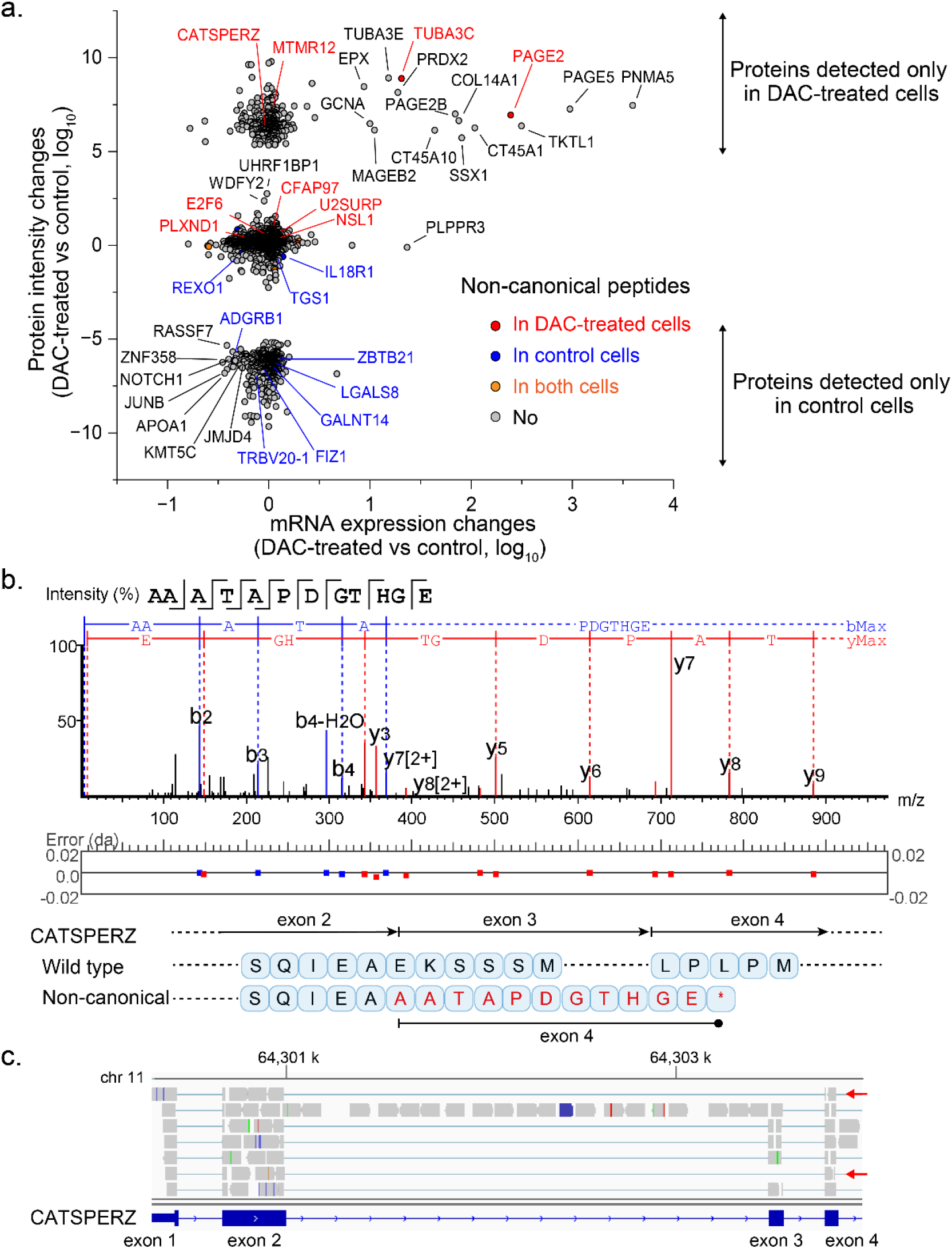
Promiscuous expression of tissue-restricted proteins, including those with non-canonical peptide sequence using ProteomeGenetator2. a. Differential protein and mRNA expression analysis reveals that DAC-treated Kasumi1 cells promiscuously express a substantial number of tissue-restricted proteins and mRNAs, some of which contain non-canonical sequence peptides. Proteins containing non-canonical sequence peptides are described as colored dots with red, blue, and orange in DAC-treated, control and both cells, respectively. b. CATSPERZ is one example, which is promiscuously expressed in DAC-treated cells and contains a non-canonical sequence peptide. c. Non-canonical sequence peptide of CATSPERZ detected in DAC-treated Kasumi1 cells is confirmed by mRNA-seq, in which exon 3 is skipped, resulting in the frameshift and truncated protein production.

## Discussion

In this work, we have laid out a computational framework for integrative proteogenomics that is modular, extensible, and accessible to users with diverse range of computational expertise. At the backbone of its versatility is the functionality to incorporate diverse collections of genetic and transcriptomic variants into an individualized and sample-specific genome reference from which sample-specific predicted protein sequences can be generated. While several contemporary proteogenomic approaches incorporate sequencing and transcriptional variation into individualized proteome databases, PG2 is the first to our knowledge that does so in an entirely harmonized manner via individualized genome against which transcriptomics analysis is performed. This uniquely enables the detection of diverse genetic, epigenetic and proteomic variation.

We have demonstrated that multiple haplotypes of such an individualized genome constitute sound, analysis-suitable genome assemblies. On a more general note, we observed that the use of an individualized genome in transcriptome sequencing and assembly can improve global accuracy and performance. That a tailored genome sequence would yield superior accuracy is perhaps unsurprising (assuming that the sequence is valid), but it remains the case that individualized genome references are not yet commonly employed in transcriptomic and proteomic studies. PG2 demonstrates that, as methods for producing individualized references continue to improve, its potential utility in such studies as an alternative to the standard genome reference should continue to be investigated.

To verify the core end-to-end functionality for integrative proteogenomics and analysis of sample-specific variation, we employed multiple validation approaches to establish the accuracy of PG2. We manually verified the presence of known genetic variants within established human cancer cell lines, and subsequently confirmed that specific alterations introduced synthetically are accurately processed by PG2. We also developed an automated framework to validate sequencing mutation propagation at genome scale, leveraging orthogonal predictions from variant annotation tools to cross reference with PG2 proteome sequences. Thus, PG2 enables variant and *de novo* transcript and protein isoform discovery without compromising performance and accuracy.

At this time, PG2 is limited by the lack of haplotype phasing between closely positioned variants, which is in part due to the limitations of short-read sequencing used in our study. However, as long-read sequencing becomes accessible, we anticipate that haplotypes will be incorporated into future versions of this method. Similarly, the current implementation of PG2 does not include a structural variant caller, which can be included in future developments. Given its scalable and modular framework, PG2 can be integrated with additional and emerging sequencing technologies, assemblers, variant callers, and mass spectral analysis algorithms. This should enable comprehensive and unbiased proteomics of diverse organisms and physiologic states.

## Conflict of interests disclosure

The authors have no competing financial interests. AK is a consultant for Novartis, Rgenta, and Blueprint.

## Author Contributions

A.K. designed the study; N.K. and Z.A. programmed and performed data analyses; S.T., P.C, and Z.S. acquired and analyzed data; A.K., N.K., and Z.A. wrote the manuscript with final editing by all authors.

## Acknowledgements

We thank members of the Kentsis Research Group for helpful feedback.

## Funding

This work was supported by the NIH R21 CA188881, R01 CA204396, P30 CA008748, Starr Cancer Consortium, St. Baldrick’s Foundation, Hyundai Hope on Wheels, Pershing Square Sohn and Mathers Foundations, Mr. William H. and Mrs. Alice Goodwin and the Commonwealth Foundation for Cancer Research and the Center for Experimental Therapeutics at MSKCC. AK is a scholar of the Leukemia and Lymphoma Society.

## Notes

https://github.com/kentsisresearchgroup/ProteomeGenerator2

https://doi.org/10.5281/zenodo.7068642

### Competing Interest Statement

The authors have declared no competing interest.

