## Supplemental Figures for "Integrative proteogenomics using ProteomeGenerator2"

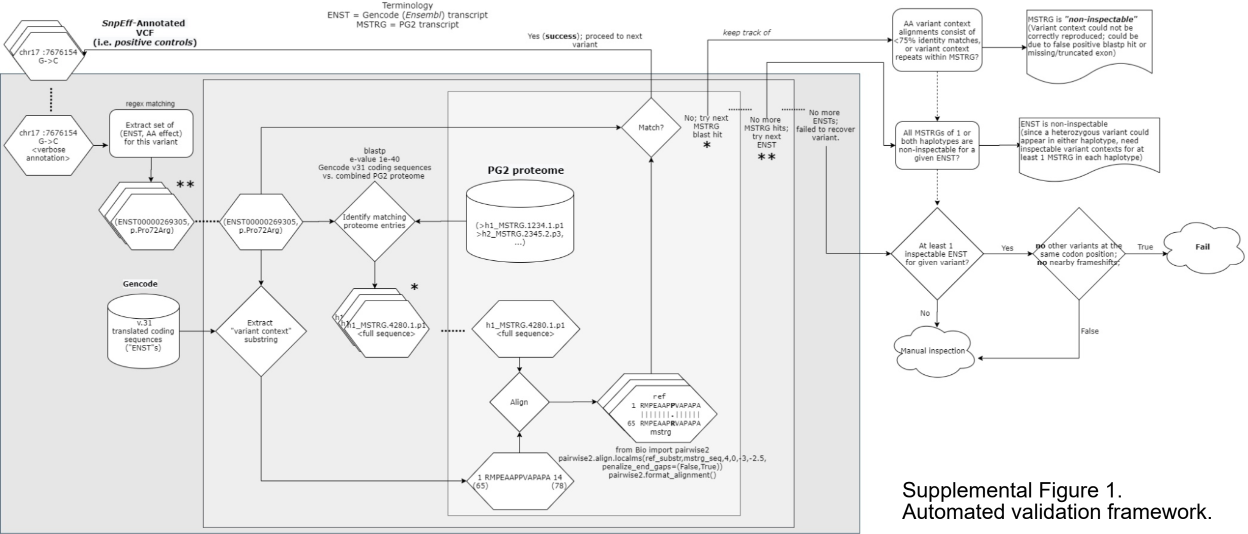


Supplemental Figure 1.
Automated validation framework.

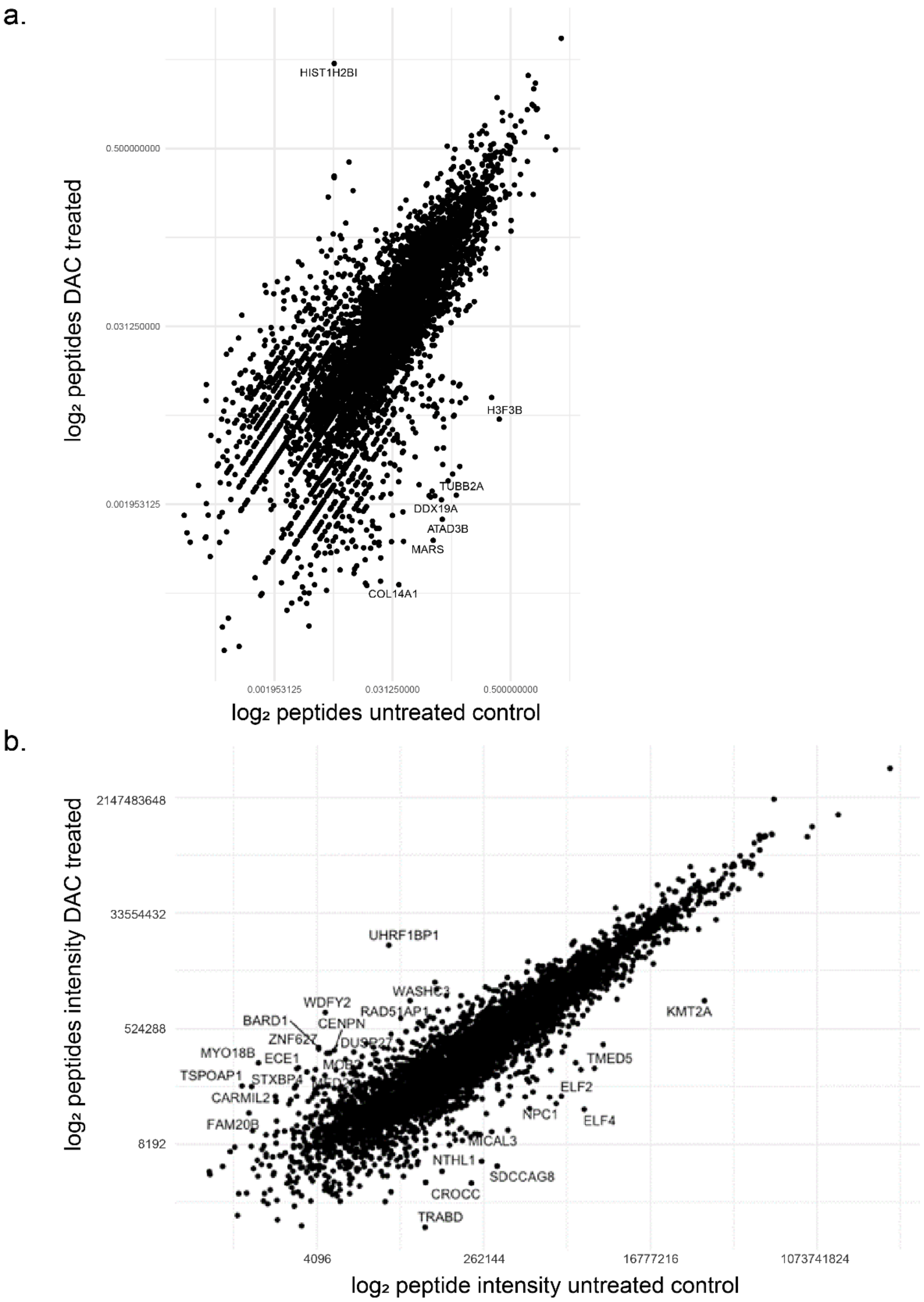


**Supplemental Figure 2. Peptide-level protein expression analysis with PG2.** Utilizing the Kasumi1 cell line dataset with paired decitabine-treated/untreated conditions, we aggregated identified enzyme-digested (including Glu-C-digested) peptides for each gene, and compared across conditions to ascertain differential protein expression. a) Volcano plot illustrating the log2-fold change in identified peptide counts per gene, across the treated/untreated conditions. To address zero values, 1 was added to every count. b) Volcano plot illustrating log2 peptide intensity detected in DAC-treated versus untreated Kasumi1 cells.
